# Mechanisms of loading and release of the 9-1-1 checkpoint clamp

**DOI:** 10.1101/2021.09.13.460164

**Authors:** Juan C. Castaneda, Marina Schrecker, Dirk Remus, Richard K. Hite

**Affiliations:** Molecular Biology Program, Memorial Sloan Kettering Cancer Center; New York, New York 10065, USA; Structural Biology Program, Memorial Sloan Kettering Cancer Center

## Abstract

5’ single-stranded/double-stranded DNA serve as loading sites for the checkpoint clamp, 9-1-1, which mediates activation of the apical checkpoint kinase, ATR^Mec1^. However, the basis for 9-1-1’s recruitment to 5’ junctions is unclear. Here, we present structures of the yeast checkpoint clamp loader, Rad24-RFC, in complex with 9-1-1 and a 5’ junction and in a post-ATP-hydrolysis state. Unexpectedly, 9-1-1 adopts both closed and planar open states in the presence of Rad24-RFC and DNA. Moreover, Rad24-RFC associates with the DNA junction in the opposite orientation of processivity clamp loaders with Rad24 exclusively coordinating the double-stranded region. ATP hydrolysis stimulates conformational changes in Rad24-RFC, leading to disengagement of DNA-loaded 9-1-1. Together, these structures explain 9-1-1’s recruitment to 5’ junctions and reveal new principles of sliding clamp loading.

## Introduction

The maintenance of genome integrity in eukaryotes is critically dependent on conserved checkpoint signaling pathways that delay the cell cycle in response to DNA damage and replication stress. Central to the induction of DNA damage and replication checkpoints is the activation of the apical checkpoint kinase, Ataxia telangiectasia and Rad3-related (ATR) in humans or Mec1 in budding yeast (*1-3*). ATR^Mec1^ activation occurs specifically on replication protein A (RPA)-coated single-stranded DNA (ssDNA) adjacent to 5’ single-stranded/double-stranded DNA (ss/dsDNA) junctions that serve as a loading site for the 9-1-1 checkpoint clamp, which mediates ATR^Mec1^ activation (*4-6*). However, how 9-1-1 is specifically targeted to 5’ ss/dsDNA junctions is not understood.

The 9-1-1 clamp is a heterotrimer composed of Rad9-Rad1-Hus1 in humans or Ddc1-Rad17-Mec3 in budding yeast that is structurally related to the canonical ring-shaped processivity clamp, proliferating cell nuclear antigen (PCNA) (*7-9*). Like all sliding clamps, 9-1-1 is loaded around DNA by an ATP-dependent clamp loader complex belonging to the AAA+ family of ATPases (*4, 10, 11*). Eukaryotes contain four related and conserved heteropentameric clamp loader complexes, termed replication factor C (RFC) and RFC-like complexes, that share four small subunits, Rfc2-5, but differ in the identity of their large subunit (*12*). 9-1-1 is loaded around DNA exclusively by the checkpoint clamp loader, Rad24-RFC in budding yeast or Rad17-RFC in humans, while RFC and other RFC-like complexes interact specifically with PCNA. Moreover, contrary to 9-1-1, PCNA is loaded around 3’ ss/dsDNA junctions. How RFC and RFC-like complexes achieve clamp and DNA substrate specificity is not understood.

In the presence of ATP, clamp loaders bind and open the clamp to allow entry of the DNA into the clamp channel, while ATP hydrolysis results in the ejection of the clamp closed around DNA (*13*). However, despite intense efforts the mechanism of clamp opening and release by eukaryotic clamp loaders is not understood. In part this is due to the lack of structural information on eukaryotic clamp loader/clamp complexes bound to DNA. Currently, the only eukaryotic clamp loader/clamp structures available are the yeast and human RFC/PCNA complexes in an inactive conformation and unbound from DNA (*14, 15*). Accordingly, it is not known how DNA is loaded onto 9-1-1, nor how it is released upon ATP hydrolysis.

To address these questions, we have reconstituted the budding yeast Rad24-RFC/9-1-1 complex bound to a DNA substrate harboring a recessed 5’ junction and examined its structure using cryogenic electron microscopy.

### Structure of Rad24-RFC:9-1-1 in complex with a DNA substrate

We expressed and purified the full-length Rad24-RFC complex composed of Rad24, Rfc2, Rfc3, Rfc4 and Rfc5, as well as the yeast 9-1-1 complex composed of Rad17, Ddc1 and Mec3, in *S. cerevisiae* cells (Fig. S1). Yeast RPA was purified after overexpression in *E. coli*, as described previously (*16*). The ATPase activity of Rad24-RFC was specifically stimulated in the presence of both DNA and 9-1-1, indicating that the proteins are functional for clamp loading (Fig. S1) (*11*). RPA was previously shown to stimulate 9-1-1 loading at 5’ junctions (*4, 5*). Therefore, to isolate the checkpoint clamp loader/clamp complex bound to 5’ junction DNA, RPA was preincubated with a DNA substrate containing a 20-base pair double-stranded DNA segment and a 50-base 3’ overhang before assembling the full Rad24-RFC:9-1-1-DNA complex in the presence of saturating ATPγS (Fig. S1). Intact complexes were separated by glycerol gradient centrifugation (Fig. S1) and then analyzed with cryogenic electron microscopy. Two distinct populations were present among the imaged particles, one that yielded a reconstruction at a resolution of 2.2 Å and a second at a resolution of 2.7 Å (Fig. S2). The high quality of the reconstructions allowed identification and modeling of Rad24-RFC, 9-1-1 and the DNA substrate (Fig. 1 and Movie S1). Despite being present in the vitrified sample, no unassigned densities are present in the map, indicating that RPA is not associated in a stable manner. In the higher resolution reconstruction, a gap of more than 30 Å is present in the 9-1-1 ring between Ddc1 and Mec3, while Ddc1 and Mec3 are in direct contact in the other reconstruction. We thus name these two states open and closed, respectively. We will first discuss the higher resolution open state.

**Fig. 1.**
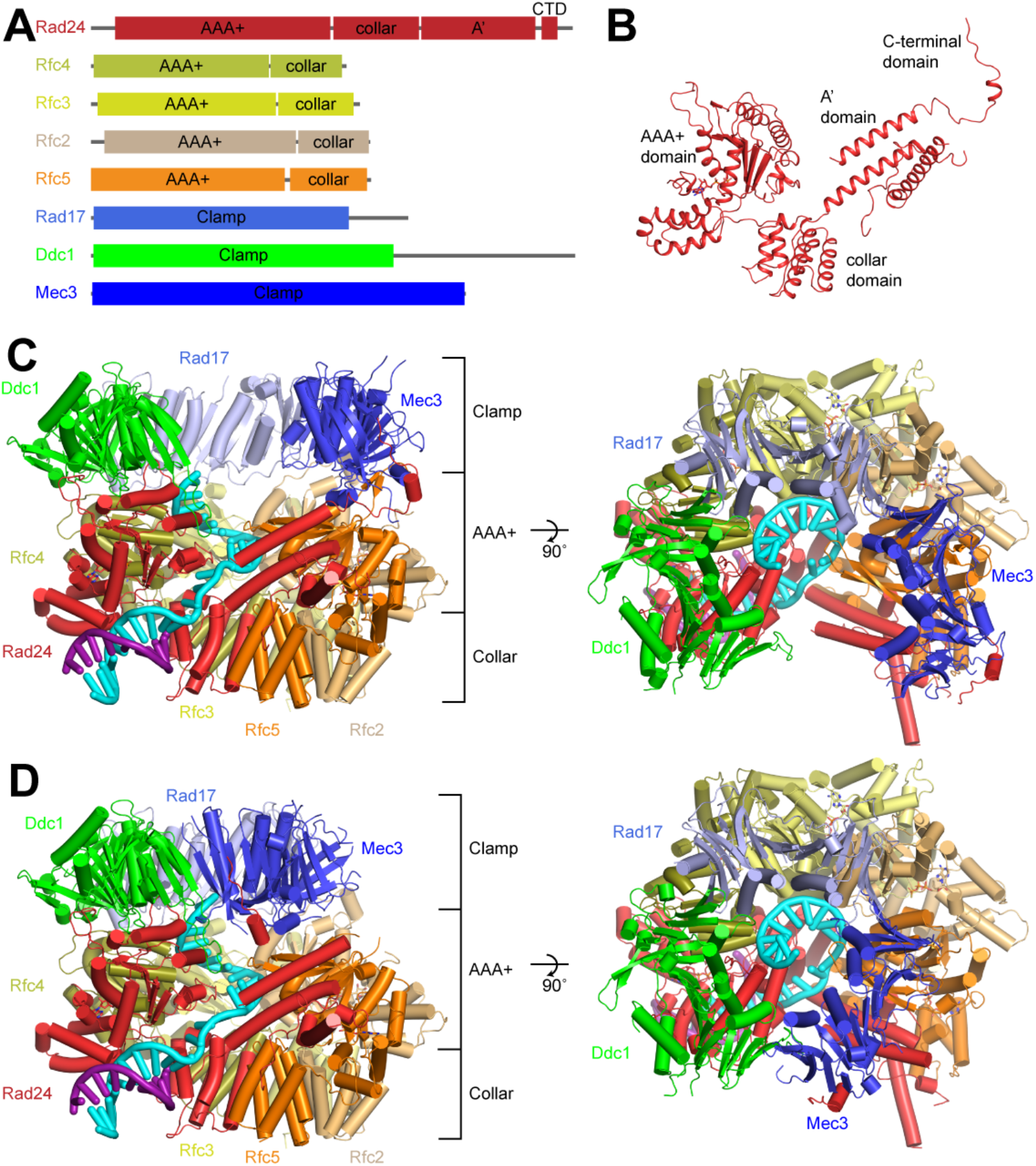
Structure of Rad24-RFC in complex with 9-1-1 and a 5’ ss/dsDNA junction. **(A)** Domain architecture of Rad24-RFC and 9-1-1. **(B)** Structure of Rad24 in Rad24-RFC:9-1-1. **(C-D)**, Structures of Rad24-RFC in complex with 9-1-1 and a 5’ junction DNA in open (**C**) and closed (**D**) states colored by subunit from side and top views. Rad24 – red, Rfc4 – brown, Rfc3 – yellow, Rfc2 – light orange, Rfc5 – orange, Ddc1 – green, Rad17 – violet, Mec3 – blue. DNA is shown as a ribbon and colored in cyan and magenta.

Viewed from the side, Rad24-RFC:9-1-1 contains three layers; a bottom layer composed of the collar domains of Rad24 and the four small RFC subunits (Rfc2-5), a middle layer composed of the AAA+ and A’ domains of Rad24 and the AAA+ domains of Rfc2-5, and a top layer composed of the 9-1-1 clamp (Rad17, Ddc1 and Mec3) (Fig. 1C). The central axis of the bottom collar domain is offset, while the top 9-1-1 layer lies flat on the middle AAA+ layer. Indeed, the AAA+ domains of all five Rad24-RFC and all three 9-1-1 subunits contribute to the clamp – clamp loader interaction surface. Rad24, Rfc3 and Rfc5, through their α4 helices, interact with the hydrophobic grooves of Ddc1, Rad17 and Mec3, respectively. Near the inter-subunit interfaces in 9-1-1, the α4 helices of Rfc4 and Rfc2 interact with Ddc1 and Rad17, respectively. Thus, the 9-1-1 clamp – clamp loader interface between Rad24-RFC and 9-1-1 is noticeably different from that resolved in structures of yeast and human RFC-PCNA complexes determined in inactive states in the absence of DNA, where the axis of the PCNA clamp is offset from the RFC clamp loader and only two of the PCNA protomers and three of the RFC subunits (Rfc1, Rfc4 and Rfc3) contribute to the clamp – clamp loader interaction surface (Fig. S3) (*14, 15, 17*). The expanded interface in Rad24-RFC:9-1-1 arises from a flattening and an expansion of the AAA+ layer. Pairwise comparisons between the subunits of Rad24-RFC and RFC in the two complexes reveal large movements of the AAA+ domains of Rad24 and Rfc4 relative to their collar domains (Fig. S3). More subtle movements occur in Rfc3 and Rfc2, while Rfc5 is unchanged. The movement of the AAA+ domain in Rad24 exposes a channel between the AAA+, collar and A’ domains of Rad24 that is contiguous with the open core of the clamp loader.

A consequence of the conformational changes in the AAA+ domains is that Rfc4, Rfc3 and Rfc2 adopt active states in which the arginine finger in the conserved SRC motif from the neighboring subunit directly participates in the coordination of the γ-phosphate of the bound ATPγS (Fig. S3). While the arginine finger of Rfc4 (Arg157) moves closer to Rad24, it remains slightly too far (4.6 Å) from the ATPγS bound to Rad24 and instead forms a salt bridge with Glu312 of Rfc4. An ADP was resolved in the nucleotide binding site of Rfc5, consistent with previous structures and its predicted structural, rather than catalytic role (*14, 15, 17*).

### DNA coordination within Rad24-RFC:9-1-1

The densities corresponding to the DNA substrate in the map were sufficiently well resolved for us to model the 10-base pair double-stranded segment preceding the junction and the 15-base single-stranded segment following the junction (Figs. 1 and 2A). The double-stranded region of the DNA protrudes out from its binding site between the AAA+ and collar domains of Rad24 (Fig. 2B). Interactions between Rad24 and phosphate groups on both strands of the dsDNA guide the junction to its binding site near the Rad24 cavity. There, the terminal base at the recessed 5’ end, C1, packs against the side of His341 and its free 5’ phosphate is coordinated by the side chains of His341, Lys345, Ser350 and His351 (Fig. 2C). Notably, while the 5’ phosphate is well coordinated, there remains sufficient room for two additional phosphate groups at the junction, potentially explaining how 9-1-1 may be loaded at Okazaki fragment 5’ ends (*18-20*). The last paired base of the junction 3’ strand, G20, forms pi-stacking interactions with the side chain of Phe340 and prevents it from base stacking with the first unpaired base, T21. The unstacked face of T21 interacts with the side chains of Tyr339, Phe340 and Phe443. Thus, Phe340, which is conserved in yeast and vertebrates (Fig. S4), appears to function analogously to the separation pin of DNA helicases (*21-23*) by separating the single-stranded and double-stranded regions of the 3’ strand, while His341 serves a similar role in stabilizing the last base of the 5’ strand.

**Fig. 2.**
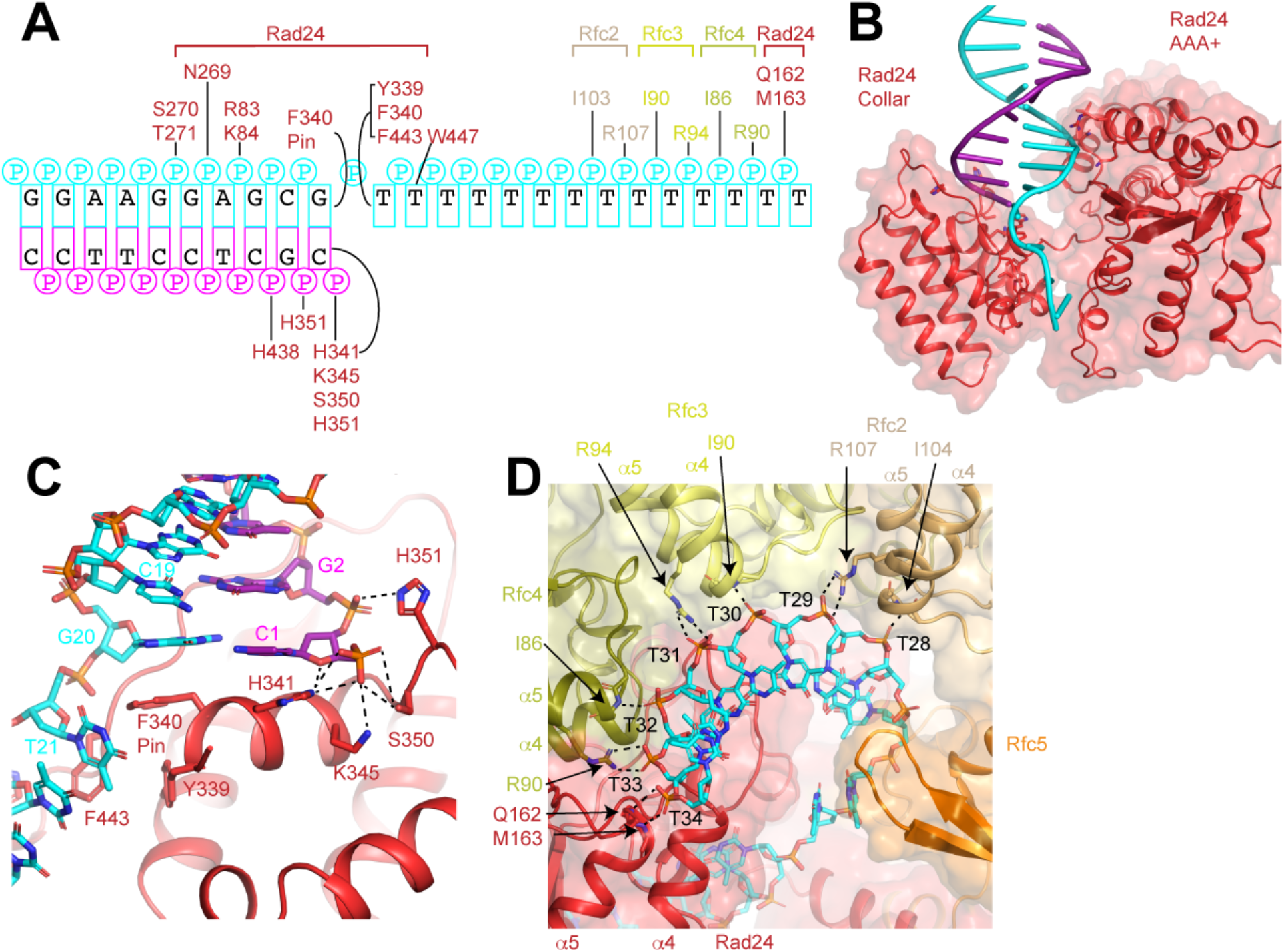
Coordination of the 5’ ss/dsDNA junction by Rad24-RFC. **(A)** Interactions between DNA and Rad24-RFC. Residues are colored as in Fig. 1. **(B)** Double-stranded DNA is coordinated by the AAA+ and collar domains of Rad24. **(C)** Coordination of the ss/dsDNA junction by Rad24. Residues that interact with the DNA are shown as sticks. Dashed lines represent polar interactions. **(D)** Coordination of ssDNA in the center of Rad24-RFC. Residues that interact with the DNA are shown as sticks. Dashed lines represent polar interactions.

From the junction, the ssDNA passes through the electropositive Rad24 channel (Fig. S4). Within the center of Rad24-RFC, the phosphate groups of the DNA form sidechain and backbone interactions with highly conserved residues on helix α5 of the RFC subunits that result in a spiral configuration closely resembling B-form DNA (Figs. 2D and S4). The phosphate of every other base (T29, T31 and T33) is coordinated by highly conserved arginine residues on Rfc2, Rfc3 and Rfc4, respectively (Figs. 2A,D and S4). Rfc5 also possesses a conserved arginine that is oriented towards the center of the clamp loader, but its side chain does not coordinate the DNA. The phosphates of the intervening bases (T28, T30, T32 and T34) form interactions with the backbone nitrogen of similarly conserved isoleucine residues at the ends of helix α5 on Rfc2, Rfc3 and Rfc4 and backbone nitrogens of Gln162 and Met163 at the ends of helix α5 of Rad24 (Figs. 2D and S4). Although these interactions establish a B-form configuration for the DNA, it is not possible for a second strand to be accommodated into the binding site. An elaborated loop in the AAA+ domain of Rad24, that is absent in Rfc1, sterically occludes double-stranded DNA from reaching into the center of Rad24-RFC (Fig. S4). Together, these interactions establish the specificity of Rad24-RFC:9-1-1 for a DNA with a 5’ junction. Moreover, while the protein-DNA interactions in the center of Rad24-RFC guide the single-stranded DNA into the center of 9-1-1, the density corresponding to the DNA within 9-1-1 is disordered in both open and closed states indicating that 9-1-1 does not make specific interactions with the DNA, consistent with its role as a sliding clamp.

### Opening of the 9-1-1 Clamp

Comparison of the open and closed states reveals that differences between the two states are largely limited to the 9-1-1 clamp (Fig. S5 and Movie S1). Rad24-RFC adopts a similar conformation in both states and the DNA follows the same path, making no discernable interactions with 9-1-1. In contrast, 9-1-1 changes from an intact ring that resembles crystal structures of human 9-1-1 and the PCNA ring in the RFC-PCNA crystal structure in the closed state (*7-9, 14*) to a horseshoe-shaped structure in the open state with a ∼30 Å gap between Mec3 and Ddc1. The ring opens through a planar, ∼40° rigid body rotation of Mec3 away from the central axis of the ring (Fig. 3A and Movie S2). Remarkably, the three 9-1-1 subunits adopt almost identical conformations and the interfacial helix α2 in Rad17, which we call the hinge helix, is the only element that undergoes significant conformational changes (Figs. 3B and S5 and Movie S2). The hinge helix, which is partially unwound in the closed state, becomes ordered in the open state and tilts ∼6 Å towards Mec3. Because the hinge helix in Rad17 is directly associated with helix α3 of Mec3, its movement must coincide with the outward movement of Mec3, thereby facilitating the opening and closing of the 9-1-1 ring.

**Fig. 3.**
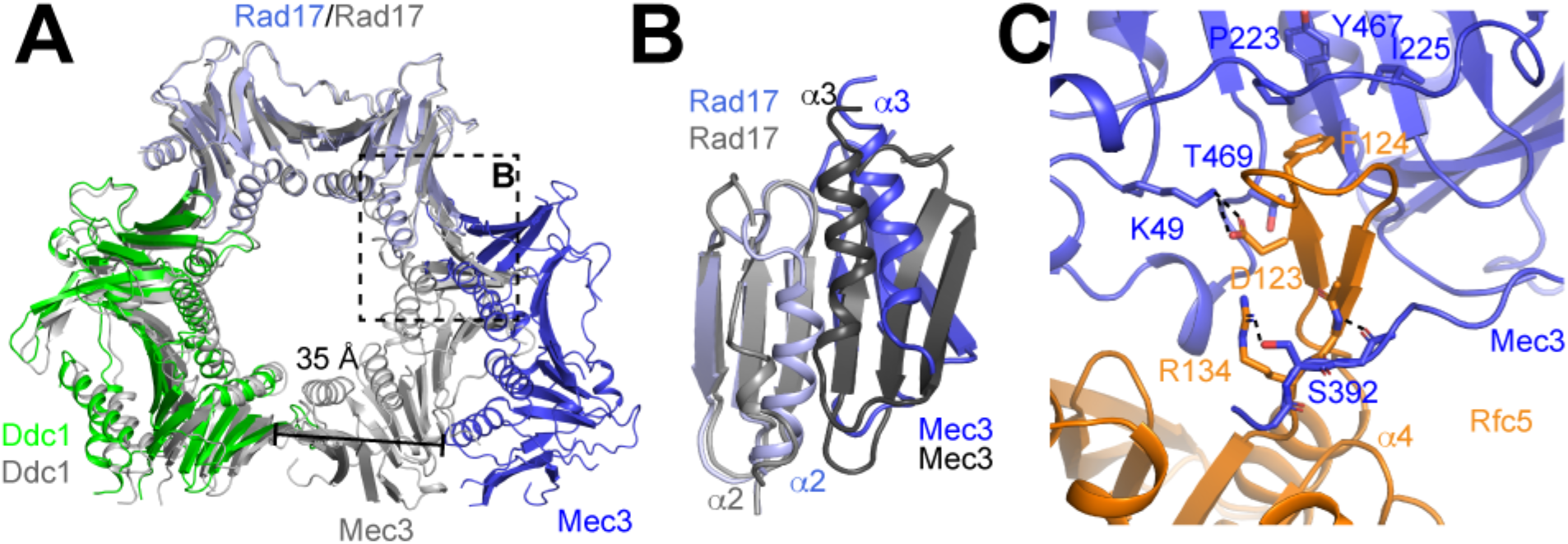
Comparison of open and closed checkpoint clamp states. **(A)** Superposition of 9-1-1 checkpoint clamp in open (colored by subunit) and closed (colored in grey) states. Dashed box is highlighted in **B. (B)** Interface between Rad17 and Mec3 in open (colored by subunit) and closed (colored in shades of grey) states. **(C)** Rfc5 binds Mec3 in the open state. Polar and ionic interactions are shown as dashed lines.

The open state of 9-1-1 is stabilized by a large interface between Mec3 and Rfc5 (1363 Å^2^), that is almost entirely absent in the closed state (80 Å^2^) (Fig. 3C). The loop following α4 in Rfc5, which is disordered in the closed state, adopts a β-hairpin structure in the open state that extends into the hydrophobic groove of Mec3. There, residues on the β-hairpin form numerous interactions with Mec3 including Asp123, which forms a salt-bridge with Lys49 of Mec3, and Phe124, which is inserted into the hydrophobic groove on the bottom of Mec3. The β-hairpin also forms backbone and side chain interactions with a loop in Mec3, which only becomes ordered upon binding to Rfc5 in the open state.

### Specificity of the checkpoint clamp loader for 9-1-1

Eukaryotic cells express 4 clamp loaders that have 4 small subunits in common, Rfc2-5, as well as one or more unique subunits that define the clamp loaders (*12*). Of the 4 clamp loaders, only Rad24-RFC (Rad17-RFC in humans) can load the 9-1-1 checkpoint clamp onto DNA. The structure of Rad24-RFC:9-1-1 reveals that Rad24 achieves specificity for 9-1-1 though an ensemble of unique interactions. In structures of DNA-free RFC-PCNA complexes, the unique subunit in the RFC complex, Rfc1, contacts PCNA exclusively through the canonical clamp loader – clamp interaction with one of the PCNA protomers through an interaction between its helix α4 and the conserved hydrophobic groove on the nearest PCNA subunit (*14, 15*). In contrast, Rad24 interacts with two 9-1-1 subunits through a combination of canonical and unique interactions (Fig. 4). Flanking the canonical interactions between helix α4 and the hydrophobic groove of Ddc1 are two loops that extend up to wrap around the central β-sheet of Ddc1 (Fig. 4A). Residues on these loops form polar and hydrophobic interactions that stabilize the interaction with Ddc1. An extended loop between β13 and β14 in Ddc1, that is not present in PCNA, binds to the side of the Rad24 AAA+ domain, further increasing the contact area. All together, these interactions establish a shared surface of 1898 Å^2^ between Rad24 and Ddc1 in open Rad24-RFC:9-1-1, more than twice that of the Rfc1-PCNA interaction in the yeast RFC:PCNA structure (928 Å^2^).

**Fig. 4.**
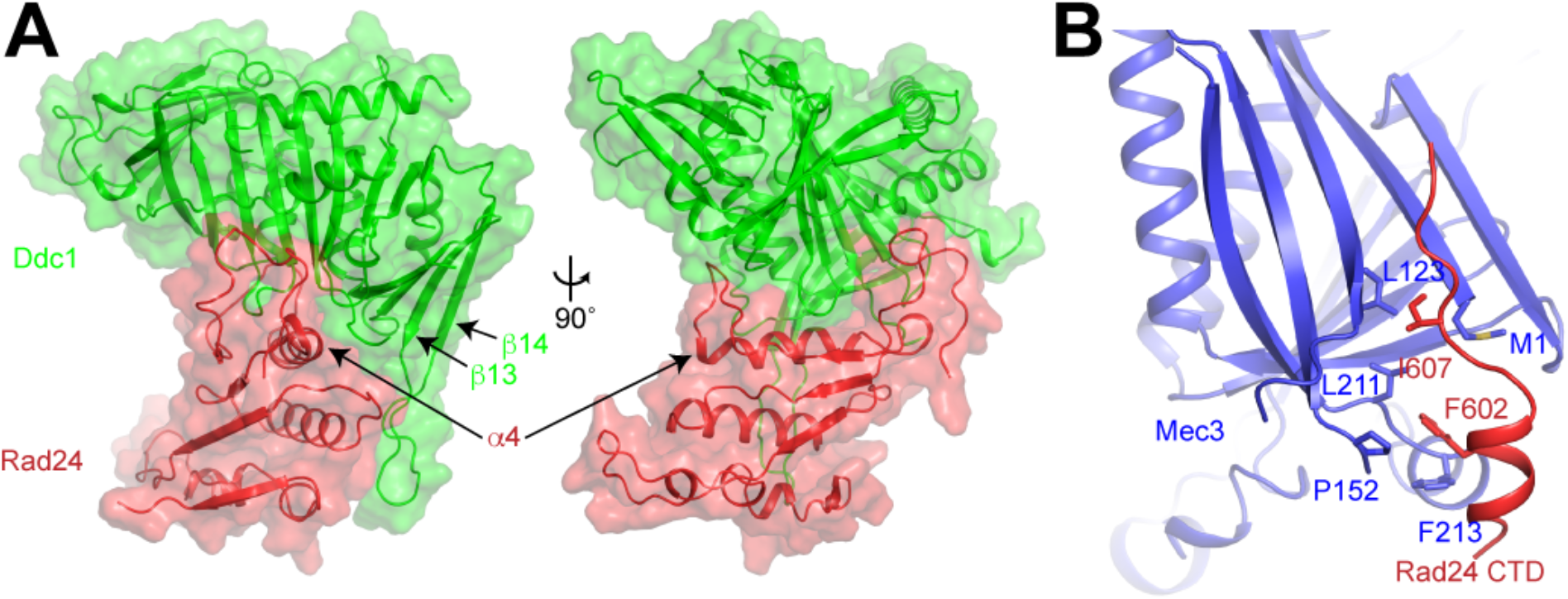
Rad24 binds to Ddc1 and Mec3. **(A)** Two views of the Rad24-Ddc1 interaction surface. **(B)** The Rad24 CTD binds to a hydrophobic pocket on the outside of Mec3.

Additional specificity for 9-1-1 arises from an interaction between Rad24 and Mec3. In both the open and closed states, a segment of the C-terminal domain of Rad24 binds to the outside of Mec3. Phe602 and Ile607 of Rad24 insert into a hydrophobic pocket on the outside of Mec3 that is lined by Met1, Leu123, Pro152, Leu211 and Phe213 (Fig 4B). The C-terminal domain is connected to the A’ domain via a flexible linker that allows the C-terminal domain to associate with Mec3 in both open and closed configurations, despite the 40 Å displacement of Mec3. Together, these interactions with Ddc1 and Mec3 allow Rad24 to establish the specificity of Rad24-RFC for 9-1-1. Moreover, as Rad24 determines the DNA specificity of Rad24-RFC:9-1-1, the interactions between Rad24 and 9-1-1 ensure that 9-1-1 is specifically loaded onto recessed 5’ DNA junctions (Fig. 2A).

### Mechanism of 9-1-1 disengagement from Rad24-RFC

Previous biochemical studies have demonstrated that 9-1-1 and the checkpoint clamp loader form a stable complex specifically in the presence of ATP or the non-hydrolyzable ATP analog, ATPγS (*4, 10, 11*). In contrast, ATP hydrolysis by the checkpoint clamp loader induces the release of DNA-loaded 9-1-1 (*4, 10, 11, 24*). To resolve a post-hydrolysis Rad24-RFC complex and gain insights into the mechanisms of clamp release, we incubated purified Rad24-RFC with ADP and analyzed its structure with cryogenic electron microscopy. We obtained a consensus reconstruction at a resolution of 2.7 Å that displays well-resolved densities for Rfc3, Rfc2 and Rfc5 (Fig. S6). The densities corresponding to the AAA+ domains of Rad24 and Rfc4 were weaker and less well resolved than those of the other subunits. Focused classification and refinement of Rad24 and Rfc4 enabled us to build and refine a model of Rad24-RFC in the presence of ADP (Fig. 5A). Comparison of Rad24-RFC_ADP_ with Rad24-RFC:9-1-1 reveals large conformational changes in the AAA+ domains, the largest of which is a 35 Å upward movement of Rad24 (Fig. S7 and Movie S3).

**Fig. 5.**
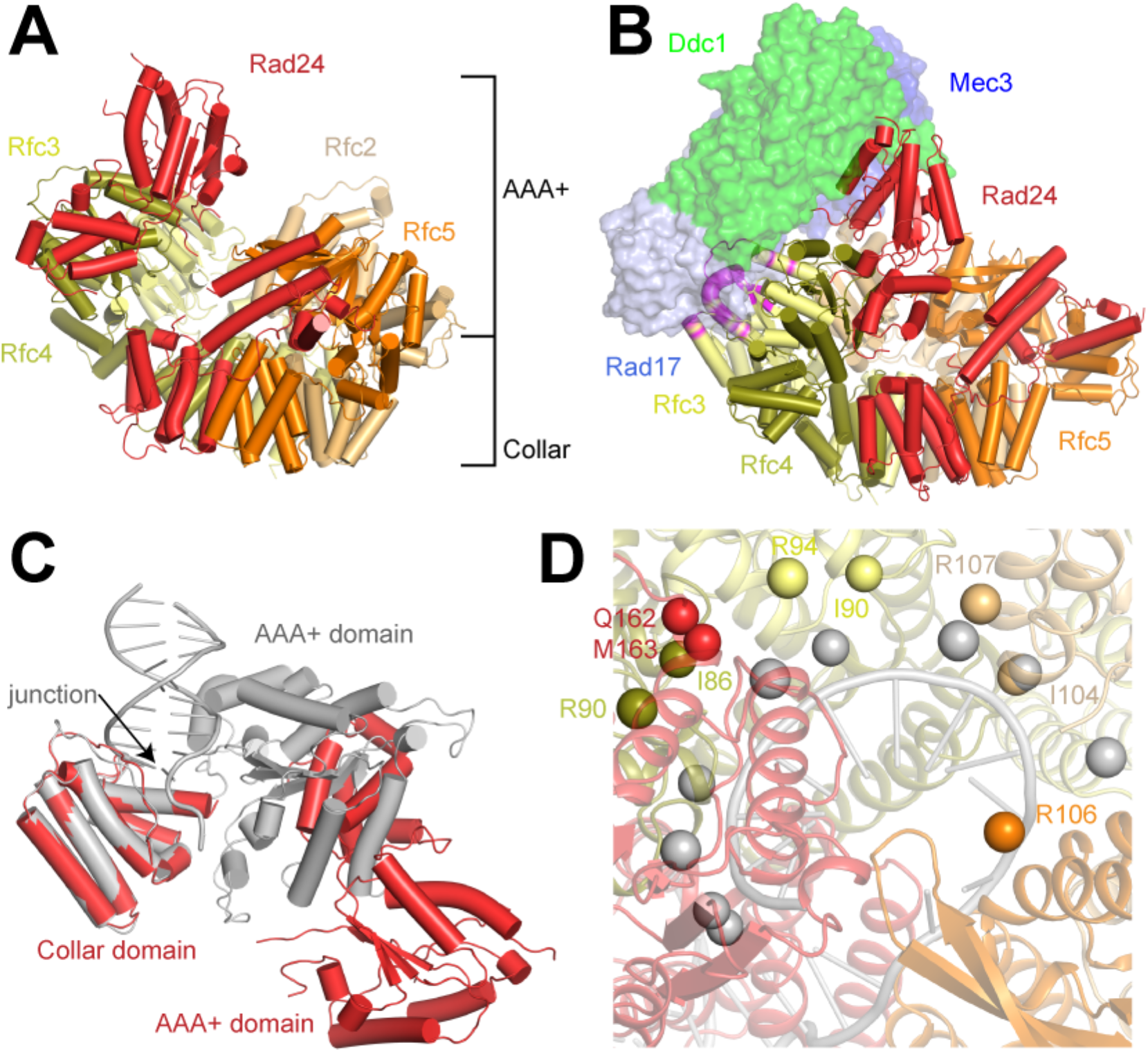
ATP hydrolysis by Rad24-RFC disrupts the 9-1-1 and DNA binding interfaces. **(A)** Structure of Rad24-RFC in the presence of ADP. Subunits are colored as in Fig. 1. **(B)** Model of Rad24-RFC_ADP_ interacting with 9-1-1. 9-1-1 is docked based on its interaction with Rad24 in Rad24-RFC:9-1-1. Rad24-RFC_ADP_ is shown as cartoon and 9-1-1 is shown as a surface. Residues on Rfc3 that sterically clash with Rad17 are colored in magenta. **(C)** Comparison of dsDNA binding site in Rad24 in Rad24-RFC_ADP_ (red) and Rad24-RFC:9-1-1 (grey). **(D)** Comparison of ssDNA interactions with the center of Rad24-RFC_ADP_ (colored by subunit) and Rad24-RFC:9-1-1 (grey). Grey spheres represent atoms that coordinate ssDNA in Rad24-RFC:9-1-1 and colored spheres represent the same atoms in Rad24-RFC_ADP_.

In Rad24-RFC:9-1-1, the γ-phosphates of the ATPγS in Rfc4, Rfc3 and Rfc2 are coordinated by the arginine fingers of helix α6 from the neighboring Rfc3, Rfc2 and Rfc5 subunits, respectively (Fig. S3). When we inspected the nucleotide densities in Rad24-RFC_ADP_, we found ADP molecules bound to Rfc4, Rfc3, Rfc2 and Rfc5 (Fig. S7). Moreover, we found that the ADP molecules in Rfc4, Rfc3 and Rfc2 are positioned ∼10 Å from the arginine finger in the neighboring subunit and that the two lobes of the AAA+ domain rock towards one another to adopt a conformation that resembles Rfc5, which binds ADP in both Rad24-RFC:9-1-1 and Rad24-RFC_ADP_ and does not undergo any conformational changes.

In contrast, despite being incubated in the presence of 0.5 mM ADP prior to vitrification, the density in the nucleotide-binding site of Rad24 is best fit by ATP (Fig. S7). Unlike the other subunits, the Rad24-RFC4 interface is only minimally altered and the bound ATP in Rad24 is still coordinated by R128 of Rfc4. As the arginine finger of Rfc4 (R157) does not adopt a catalytic conformation in Rad24-RFC:9-1-1, the bound nucleotide may serve a structural role in Rad24, stabilizing the interface with Rfc4, which may explain why the mutation of the Rad24 Walker A lysine residue (K157E) disrupts Rad24-RFC function (*24, 25*) (Fig. S3).

How does nucleotide hydrolysis by Rfc4, Rfc3 and Rfc2 alter the relationship between Rad24-RFC and DNA-bound 9-1-1? The reorganization of the AAA+ layer alters the relative position of the AAA+ domains, rendering it impossible to maintain interactions between 9-1-1 and all five Rad24-RFC subunits. Indeed, Rfc4 is the only other subunit that can favorably interact with 9-1-1 when either the open or closed 9-1-1 ring is docked onto Rad24-RFC_ADP_ based on the Rad24-Ddc1 interaction resolved in Rad24-RFC:9-1-1 (Fig. 5B). Rfc2 and Rfc5 do not contact 9-1-1 in the docking models, while Rfc3 clashes extensively with Rad17. Thus, nucleotide hydrolysis disrupts the association between Rad24-RFC and 9-1-1.

Nucleotide hydrolysis also disrupts the interactions between Rad24-RFC and the DNA. In Rad24-RFC:9-1-1, the AAA+ and collar domains together establish the double-stranded DNA binding site (Fig. 2B). In Rad24-RFC_ADP_, the AAA+ domain moves ∼35 Å away, separating the two halves of the binding site and releasing the bound DNA (Fig. 5C). Moreover, the movement of the Rad24 AAA+ domain causes a reorganization of the AAA+-collar linker, which includes key residues that contribute to the ss/dsDNA junction binding site. The rearrangement of the AAA+ domains also disrupts interactions with the ssDNA in the center of Rad24-RFC (Fig. 5D). In Rad24-RFC:911, the ssDNA is coordinated by backbone and side chain interactions with Rfc2, Rfc3, Rfc4 and Rad24. These residues are all moved with respect to one another and cannot stably bind the DNA in Rad24-RFC_ADP_.

Together, these changes induced by ATP hydrolysis yield a conformation for Rad24-RFC that is incompatible with binding to 9-1-1 or the DNA, thus suggesting a mechanism for 9-1-1 release from Rad24-RFC.

### Model for checkpoint clamp loading and release

Our structural data allow us to posit a model for the loading of 9-1-1 at 5’ ss/dsDNA junctions and for the ATP hydrolysis-induced release of the DNA-loaded 9-1-1 clamp (Fig. 6). In this model, ATP binding to Rad24-RFC promotes a partial engagement of 9-1-1 by Rad24-RFC, analogous to the RFC-PCNA structure obtained in the absence of DNA (*14*). Although Rfc2-5 are principally capable of interacting with either 9-1-1 or PCNA, the large interaction surface between Rad24 and Ddc1/Mec3 confers selectivity to the 9-1-1/Rad24-RFC interaction. Complete engagement of 9-1-1 by Rad24-RFC requires the binding of Rad24-RFC to DNA, which locks a flexible module formed by the Rfc4 and Rad24 AAA+ domains in a stable conformation to form a dsDNA binding channel between the AAA+ and collar domains of Rad24. In this configuration, Rad24-RFC and 9-1-1 can adopt a co-planar clamp loader/clamp conformation that allows the five clamp loader subunits to intimately engage all three 9-1-1 subunits.

**Fig. 6.**
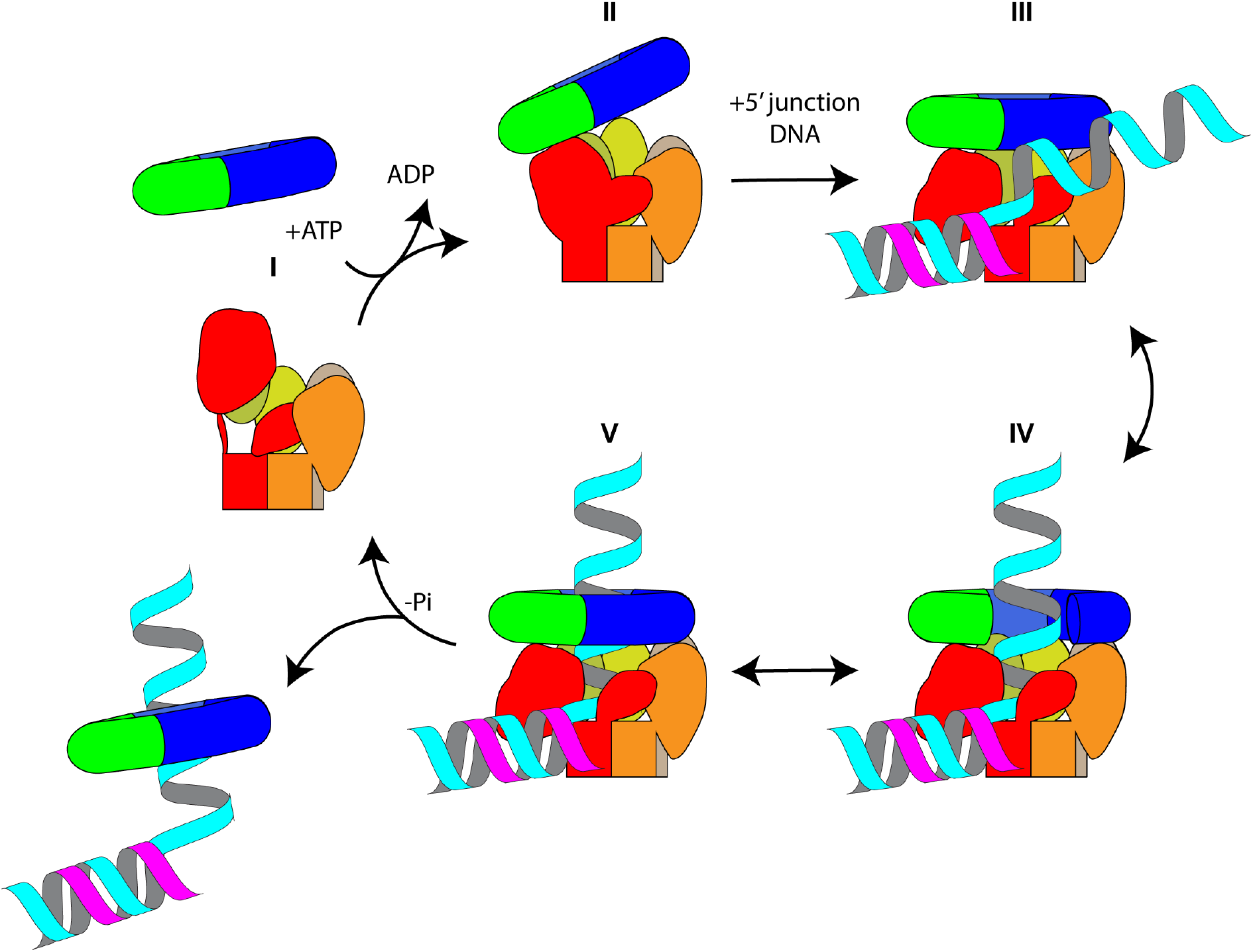
Model for loading and release of the 9-1-1 checkpoint clamp by Rad24-RFC. (I) In the presence of ADP, Rad24 adopts an up conformation. (II) Exchange of ADP with ATP in Rfc4, Rfc3 and Rfc2, leads to conformational changes in the AAA+ domains that establish a platform for off-axis clamp binding. (III) Upon recognition of a 5’ DNA junction by Rad24-Rfc:9-1-1, AAA+ domains rearrange and Rad24 adopts a down conformation, enabling on-axis interactions between the clamp and all 5 AAA+. (IV) Clamp loader opens 9-1-1 ring and enables coordination of the ssDNA within the clamp loader chamber. (V) ATP hydrolysis in Rfc4, Rfc3 and Rfc2 induces release of DNA-bound 9-1-1.

While Rad24 is well positioned to bind dsDNA, a protrusion in the Rad24 AAA+ domain limits the central Rad24-RFC chamber to ssDNA binding, thus orienting Rad24-RFC at 5’ ss/dsDNA junctions in reverse orientation relative to RFC at 3’ ss/dsDNA junctions (*13*). In addition, the specific coordination of the 5’ ss/dsDNA junction by Rad24 and a base stacking interaction by the Rad24 pin residue, F340, locks Rad24-RFC in position at the junction, placing 9-1-1 over the ssDNA region of the junction. Thus, contrary to PCNA, 9-1-1 is initially loaded on ssDNA, although it may slide over dsDNA at the junction upon release from the clamp loader (*4, 10, 11*).

The 9-1-1 clamp can engage DNA- and ATP-bound Rad24-RFC in an open or closed configuration. 9-1-1 opening occurs specifically at the Ddc1-Mec3 interface near the gap formed between the Rad24 and Rfc5. While the open and closed states of 9-1-1 bound to Rad24-RFC may be energetically similar, the open state is stabilized by induced specific interactions between Rfc5 and Mec3. In contrast, the interaction surfaces between Ddc1/Rad17 and Rad24-RFC remain largely unchanged in both states and may thus act as a stator to facilitate the Mec3 swinging movement during clamp opening. Clamp opening allows subsequent entry of the ssDNA into the 9-1-1 channel. Remarkably, at > 30 Å, the gap formed between Ddc1 and Mec3 greatly exceeds the gap size required for ssDNA entry, and also greatly exceeds the gap size reported for other sliding clamps (*17, 26, 27*). Why the clamp needs to be opened to such an extent is not clear, but it is noteworthy that ssDNA at the junction is likely to be bound by RPA, which promotes the loading of the checkpoint clamp at 5’ junctions (*4, 5*). Extended clamp opening may thus help negotiate the RPA-ssDNA filament during 9-1-1 loading.

Following the placement of 9-1-1 on DNA, ATP hydrolysis by the checkpoint clamp loader triggers 9-1-1 release (*4, 10, 11, 24*). This release is induced by steric clashes between the planar 9-1-1 ring and ADP-bound Rad24-RFC. The structural changes in Rad24-RFC upon ATP hydrolysis furthermore disrupt Rad24-RFC-DNA interactions, providing a mechanism for the release of Rad24-RFC from the DNA junction following 9-1-1 loading. Thus, while previous models have implicated the non-planar conformation of clamp-loaders in clamp opening (*13*), our data suggests that the non-planar conformation of Rad24-RFC is important for 9-1-1 release, while the planar Rad24-RFC conformation stabilizes the open state of 9-1-1. Whether this unexpected clamp opening and release mechanism is specific for the eukaryotic checkpoint clamp loader system or can be generalized to RFC and other RFC-like complexes will be the subject of future studies.

## Materials and Methods

### Protein expression and purification

Plasmids used in this study are summarized in Table 1. Proteins used in this study and a summary of each protein’s purification steps are listed in Supplementary Table 2. RPA was purified as described previously (*16*).

Yeast strains used in this study are listed in Table 3. Yeast cells carrying galactose-inducible expression constructs were grown at 30°C in YP-GL (YP + 2% glycerol/2% lactic acid) to a density of 2-4 × 10^7^ cells/ml. Galactose was then added to 2% and cell growth continued for 2 or 4 hours for 9-1-1 and Rad24^Rfc^, respectively. Cells were harvested by centrifugation and washed once with 1 M sorbitol/25 mM Hepes-KOH pH 7.6 followed by a second wash with buffers as indicated. Washed cells were resuspended in 0.5 volumes of respective buffers as indicated and frozen dropwise in liquid nitrogen; the resulting popcorn was stored at -80°C. Frozen popcorn was crushed in a freezer mill (SPEX SamplePrep 6875D Freezer/Mill) for 6 cycles of 2 min at a rate of 15 impacts per second. Extracts were clarified by centrifugation at 40,000 rpm for 1 hr (T647.5 rotor). All proteins were stored in aliquots at -80 °C.

#### 9-1-1

YJC8 cells were washed with buffer A (50 mM Hepes-KOH pH 7.6/150 mM KCl / 1 mM EDTA 1 mM EGTA/10% glycerol/0.02% NP40S/1 mM DTT) and resuspended in ½ volume of buffer A supplemented with protease inhibitors for preparation of cell popcorn. Cell powder was thawed on ice and resuspended in one volume of buffer A for clarification of the extract by ultracentrifugation. Clarified extract was passed over M2-agarose anti-FLAG beads at 4°C using Bio-Rad EconoColumn at a flow rate of 3 mL/min. Beads were washed with 6 CV of buffer A. Bound protein was eluted with 1 CV of buffer A/FLAG peptide (0.5 mg/mL), followed by 2 CVs of buffer A/FLAG peptide (0.25 mg/mL), and then 2 CVs of buffer A. Eluates were pooled and fractionated on a Mono Q column using a salt gradient of 0.15 – 1 M KCl over 20 CVs. Peak fractions from the Mono Q step were pooled and fractionated on a 24 mL Superdex 200 Increase gel filtration column equilibrated in 25 mM Hepes-KOH pH 7.6/300 mM KOAc /1 mM EDTA/10% glycerol/1mM DTT. Peak fractions from S200 step were pooled and concentrated on Amicon Ultracel 30K 0.5 mL filter units.

#### Rad24^Rfc^

YJC3 cells were washed with buffer A (50 mM Hepes-KOH pH 7.6/150 mM KCl/1 mM EDTA/1 mM EGTA/10% glycerol/0.02% NP40S/1 mM DTT) and resuspended in ½ volume of buffer A supplemented with protease inhibitors for preparation of cell popcorn. Cell powder was thawed on ice and resuspended in one volume of buffer A for clarification of the extract by ultracentrifugation. Clarified extract was passed over M2-agarose anti-FLAG beads at 4°C using Bio-Rad EconoColumn at a flow rate of 3 mL/min. Beads were washed with 6 CV of buffer A. Bound protein was eluted with 1 CV of buffer A/FLAG peptide (0.5 mg/mL), followed by 2 CVs of buffer A/FLAG peptide (0.25 mg/mL), and then 2 CVs of buffer A. Eluates were pooled together and fractionated on Mono Q column with a salt gradient of 0.15 – 1 M KCl over 20 CVs. Peak fractions from the Mono Q step were pooled and dialyzed into 25 mM Hepes-KOH pH 7.6/300 mM KOAc/1 mM EDTA/10% glycerol/1mM DTT. Dialyzed fraction was concentrated on Amicon Ultracel 30K 0.5 mL filter units.

#### Preparation of DNA template

Oligos were purchased from IDT. DR2636: 5’-GGACGAGTCAGGAAGGAGCGTTTTTTTTTTTTTTTTTTTTTTTTTTTTTTTTTTTTTTTTTTTTTTTTTT-3’; DR2628: 5’-phosphate/CGCTCCTTCCTGACTCGTCC-3’. DR2636 was PAGE-purified prior to template generation. The DNA oligos were annealed by mixing to a final concentration of 10 μM in 1x TE pH 7.5/150 mM NaCl, heat denaturation at 95°C, and gradually decreasing the temperature from 95°C to 10°C in steps of 1°C per minute using a thermo cycler.

#### ATPase Assays

Reactions were carried out in buffer containing 15 mM Hepes-KOH pH 7.6/160 mM KOAc/5 mM NaCl/7 mM Mg(OAc)_2_/0.6 mM EDTA/6% glycerol/0.6 mM DTT/25 μM ATP (incl. [α-^32^P]ATP). Rad24-RFC, 9-1-1, and DNA substrate were included at 30 nM, 100 nM, and 600 nM, respectively, as indicated. Reactions were incubated at 30°C and at the indicated times 2 μL were spotted on TLC PEI Cellulose F (Millipore) to stop the reaction. The TLC plates were developed in 0.6 M sodium phosphate buffer, scanned on a Typhoon FLA 7000 phosphorimager, and ATP hydrolysis quantified using ImageJ.

#### Preparation of samples for cryo-EM grids

All incubation steps were carried out at 30°C. First, 9-1-1 (4.11 μM) and Rad24-RFC (3.78 μM) were mixed for 30 minutes in a 51.1 μL reaction volume in a buffer containing 25 mM Hepes-KOH pH 7.6/300 mM KOAc/10% glycerol/1 mM DTT/1.94 mM ATPγS/9.7 mM Mg(OAc)_2_. Subsequently, RPA (30 nM) and DNA (1.5 μM) were added to the reaction, the reaction volume was increased a final volume of 100 μL with reaction buffer (25 mM Hepes-KOH pH 7.6/300 mM KOAc/7 mM Mg(OAc)_2_/5% glycerol/0.02% NP40S/1 mM DTT/12.5 mM Mg(OAc)_2_), and incubation continued for 30 min. The reaction was layered onto a 4 mL 10 – 35% glycerol gradient containing 25 mM Hepes-KOH pH 7.6/300 mM KOAc/7 mM Mg(OAc)_2_/0.02% NP40S/1 mM DTT. Gradients were centrifuged at 45,000 rpm for 6 hours at 4°C in a Thermo Scientific AH-650 swing bucket rotor. 200 μL fractions were manually collected from the top of the gradient and 10 μL of each fraction analyzed by SDS-PAGE stained with SilverQuest Staining Kit (Invitrogen). Another 10 μL of each fraction were subjected to DNA quantification using a QuBit 3.0 Fluorometer with QuBit dsDNA HS Assay kit (Q32851). Peak fractions were pooled and dialyzed against buffer containing 25 mM Hepes-KOH pH 7.6/300 mM KOAc/7 mM Mg(OAc)_2_. Dialyzed sample was concentrated via centrifugation in Amicon Ultracel 30K 0.5 mL filter units (UFC503096) and the protein concentration of the final sample determined by SDS-PAGE and Coomassie stain by comparing with a known protein standard.

#### Cryo-EM sample preparation and data acquisition

For the Rad24-Rfc:9-1-1 complex from *S. cerevisiae*, 3.5 μl of purified protein at a concentration of 1.0 mg/ml was applied to Graphene Oxide Au 400 mesh QUANTIFOIL R1.2/1.3 holey carbon grids (Quantifoil), and then plunged into liquid nitrogen-cooled liquid ethane with a FEI Vitrobot Mark IV (FEI Thermo Fisher). The sample was frozen at 4°C with 100% humidity, using blotting times between 30 and 60 s and a waiting time of 30 s. For Rad24-Rfc, 3.5 μl of purified protein at a concentration of 4.5 mg/ml was supplemented with 0.5 mM ADP followed the same freezing conditions as for the Rad24-Rfc:9-1-1 complex. Grids were transferred to a 300 keV FEI Titan Krios microscopy equipped with a K3 summit direct electron detector (Gatan). Images were recorded with SerialEM(*28*) in super-resolution mode at 29,000x, corresponding to super-resolution pixel size of 0.413 Å. Dose rate was 15 electrons/pixel/s, and defocus range was −0.5 to −2.0 µm. Images were recorded for 3 s with 0.05 s subframes (total 60 subframes), corresponding to a total dose of 66 electrons/Å^2^.

#### Cryo-EM processing

60-frame super-resolution movies (0.413 Å/pixel) of Rad24-Rfc:9-1-1 were gain corrected, Fourier cropped by two (0.826 Å/pixel) and aligned using whole-frame and local motion correction algorithms by cryoSPARC v3.2.0(*29*). Blob-based autopicking in cryoSPARC was implemented to select initial particle images with 4,117,022 particles. Several rounds of two-dimensional classification were performed and the best 2D classes were manually selected for the initial 3D model generation using the *ab initio* algorithm in cryoSPARC. False-positive selections and contaminants were excluded through iterative rounds of heterogeneous classification using the model generated from the *ab initio* algorithm, resulting in a stack of 938,420 particles for the open state and 237,512 particles for the closed state. After particle polishing in Relion 3.1.2(*30*) and local CTF estimation and higher order aberration correction in cryoSPARC v3.2.0, a reconstruction was determined at estimated resolutions of 2.20 Å for the open state and of 2.72 Å for the closed state by non-uniform refinement in cryoSPARC v3.2.0(*31*). A focused refinement of the open state particles using a mask encompassing the Mec3 yielded a map with improved interpretability for Mec3 with an estimated resolution of 2.43 Å. The reconstructions were further improved by employing density modification on the two unfiltered half-maps with a soft mask in Phenix(*32*).

60-frame super-resolution movies (0.413 Å/pixel) of Rad24-Rfc were gain corrected, Fourier cropped by two (0.826 Å/pixel) and aligned using whole-frame and local motion correction algorithms by cryoSPARC v3.2.0 (*29*). 4,432,421 particles were picked using blob-based autopicking in cryoSPARC. Several rounds of two-dimensional classification were performed and the best 2D classes were manually selected and used to generate initial 3D model using the ab initio algorithm in cryoSPARC. False-positive selections and contaminants were excluded through iterative rounds of heterogeneous classification using the model generated from the ab initio algorithm, resulting in a stack of 183,334 particles. After particle polishing in Relion(*30*) and local CTF estimation and higher order aberration correction in cryoSPARC v3.2.0(*29*), a reconstruction was determined at resolution of 2.73 Å by non-uniform refinement in cryoSPARC v3.2.0. Focused three-dimensional classification in Relion identified a sub-population of 81,834 particles that displayed similar densities for the AAA+ domains of Rad24 and Rfc4. A consensus reconstruction of these particles achieved an estimated resolution of 2.90 Å and focused refinement using a soft mask encompassing the AAA+ domains of Rad24 and Rfc4 yielded a map with improved interpretability with an estimated resolution of 3.89 Å. The reconstructions were further improved by employing density modification on the two unfiltered half-maps with a soft mask in Phenix(*33*).

#### Model building and refinement

The structures of the yeast RFC-PCNA complex (PDB: 1SXJ)(*14*) and human 9-1-1 complex (PDB: 3A1J)(*8*) were manually docked into the open structure of Rad24-RFC:9-1-1 in UCSF Chimera(*34*). The models were manually rebuilt to fit the density and sequence in COOT(*35*) using the consensus and Mec3-focused refinement maps. The models were initially refined in ISOLDE(*36*) to correct geometric errors before several cycles of manual rebuilding in COOT and real space refinement in Phenix(*33*) against the consensus open state map.

The refined open state model was manually docked into closed state map and manually rebuilt to fit the density in COOT. The final model was subjected to real space refinement in Phenix against the closed state map.

The refined open state model was manually docked into the Rad24-RFC_ADP_ map in Chimera. The model was manually rebuilt to fit the density of the consensus and Rad24-focused maps in COOT. The final model was subjected to real space refinement in Phenix against the consensus Rad24-RFC_ADP_ map.

Figures were prepared using PyMol (www.pymol.org), UCSF Chimera(*34*) and UCSF ChimeraX(*37*).

## Acknowledgments

We thank M de la Cruz at the MSKCC Richard Rifkind Center for cryo-EM for assistance with data collection and the MSKCC HPC group for assistance with data processing.

## Funding

NIH-NCI Cancer Center Support Grant P30 CA008748

NIGMS R01-GM107239 (DR)

NIGMS R01-GM127428 (DR)

Walter Benjamin Fellow of the Deutsche Forschungsgemeinschaft (MS)

Josie Robertson Investigators Program (RKH)

## Author contributions

Conceptualization: JCC, MS, DR, RH

Methodology: JCC, MS, DR, RH

Analysis: JCC, MS, DR, RH

Funding acquisition: MS, DR, RH

Supervision: DR, RH

Writing – original draft: JCC, MS, DR, RH H

Writing – review & editing: JCC, MS, DR, RH

## Data and materials availability

Cryo-EM maps and atomic coordinates have been deposited with the EMDB and PDB under accession codes EMDB-XXXXX and PDB XXXX for Rad24-RFC:9-1-1 in the open state, codes EMDB-XXXXX and PDB XXXX for Rad24-RFC:9-1-1 in the closed state, and codes EMDB-XXXXX and PDB XXXX for Rad24-RFC_ADP_.

