## Supplementary Figures for "Mechanisms of loading and release of the 9-1-1 checkpoint clamp"

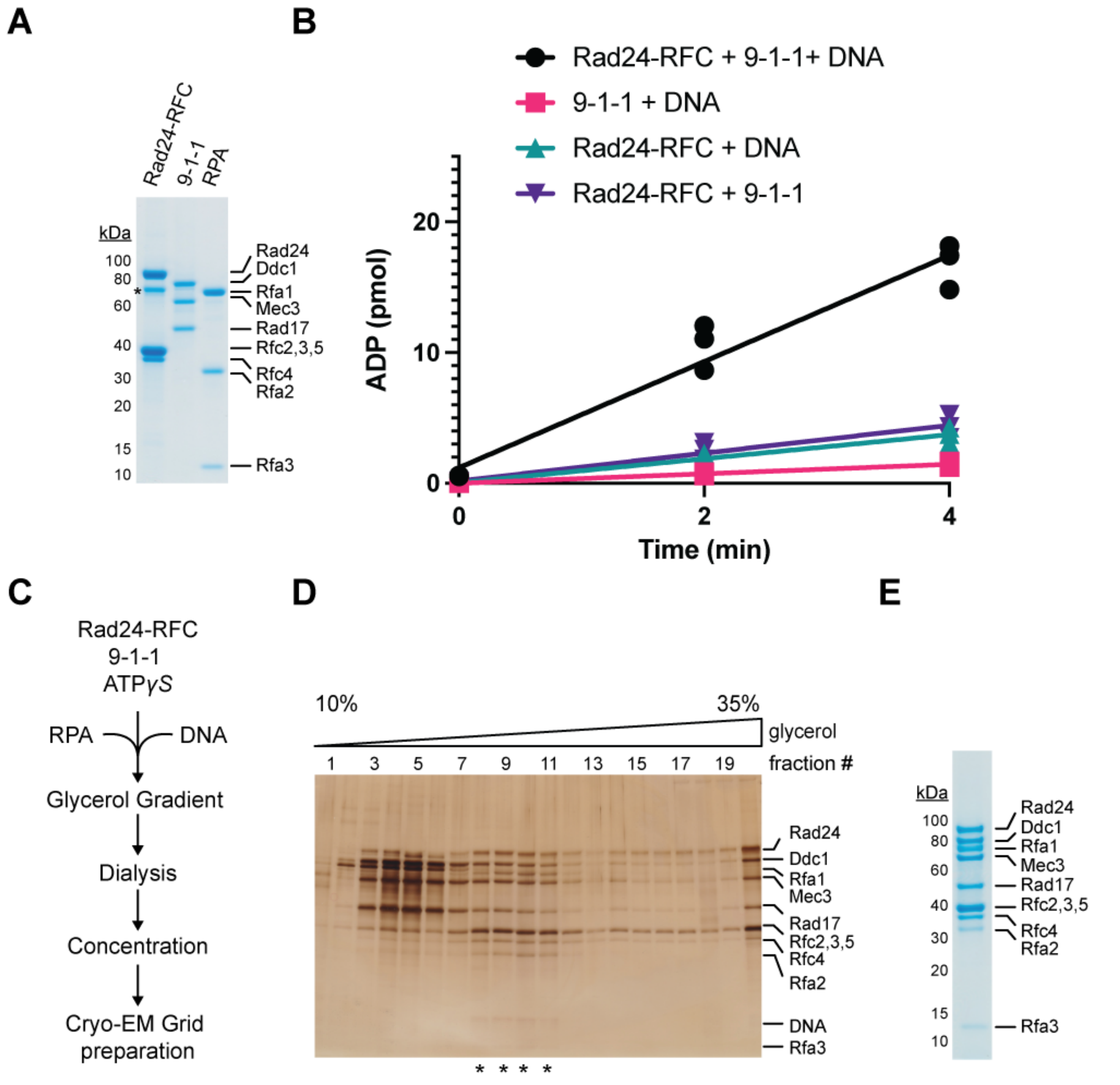


**Fig S1. Purification and analysis of Rad24-RFC:9-1-1. (A)** Representative Coomassie-stained SDS-PAGE analysis of purified *S. cerevisiae* Rad24-RFC, 9-1-1 and RPA. The * corresponds to truncated form of Rad24. **(B)** ATPase activity of Rad24-RFC in the presence of 9-1-1 and DNA (black circles), Rad24-RFC in the presence of DNA (teal triangles), Rad24-RFC in the presence of 9-1-1 (purple triangles) and 9-1-1 and DNA (pink squares). Experiments are shown in triplicate. **(C)** Schematic for the assembly of Rad24-RFC:9-1-1:DNA in the presence of ATPγS. **(D)** Representative silver-stained SDS-PAGE analysis of Rad24-RFC:9-1-1 fractions following glycerol gradient (10-35%) centrifugation. Fractions pooled for Cryo-EM analysis are denoted with *. **(E)** Representative Coomassie-stained SDS-PAGE analysis of purified *S. cerevisiae* Rad24-RFC:9-1-1:DNA.

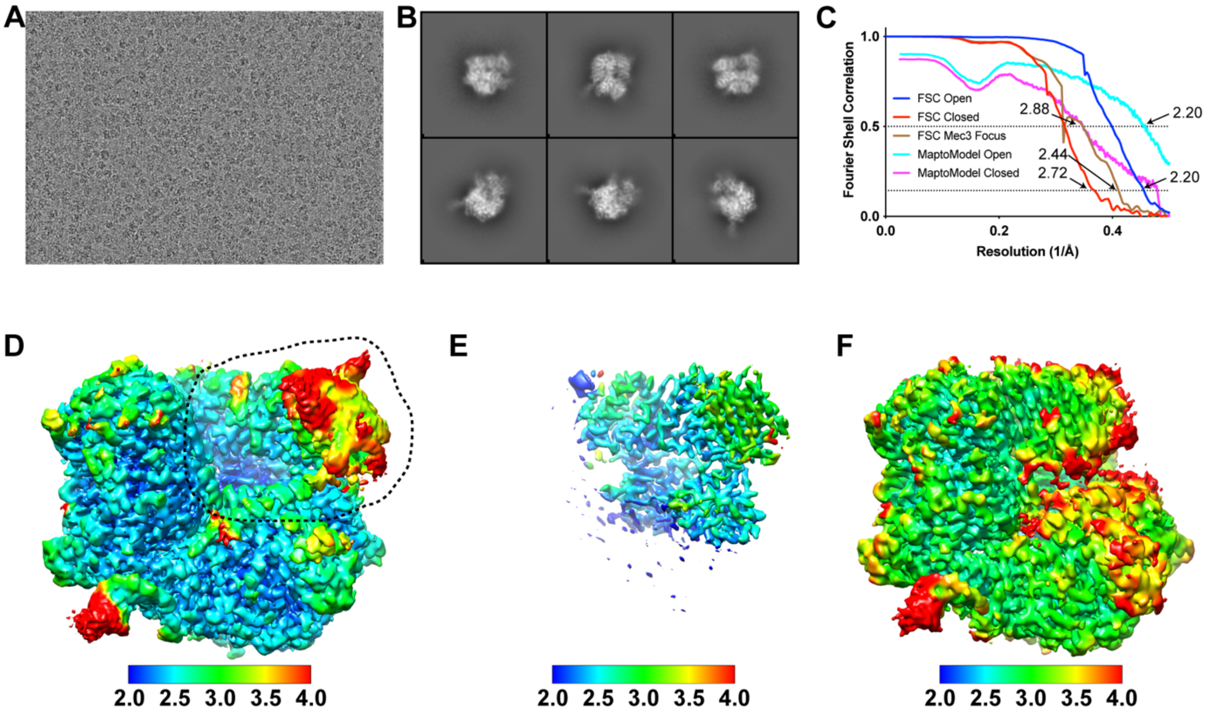


**Fig S2. Validation of Rad24-RFC:9-1-1 structures. (A)** Representative cryo-EM image of vitrified Rad24-RFC:9-1-1:DNA. **(B)** Representative two-dimensional averages of Rad24-RFC:9-1-1. **(C)** Plot of Fourier shell correlations between two independent open state half-maps (blue), two independent closed state half-maps (red), two independent Mec3 focus half-maps (brown), the open state map and open state atomic model (cyan), and the closed state map and closed state atomic model (magenta). **(D-F)** Cryo-EM density maps of open (**D**), Mec3-focused refined (**E**) and closed (**F**) maps colored by local resolution. Region included in the mask for the Mec3 focused refinement (**E**) is denoted by the dashed line in **D**.

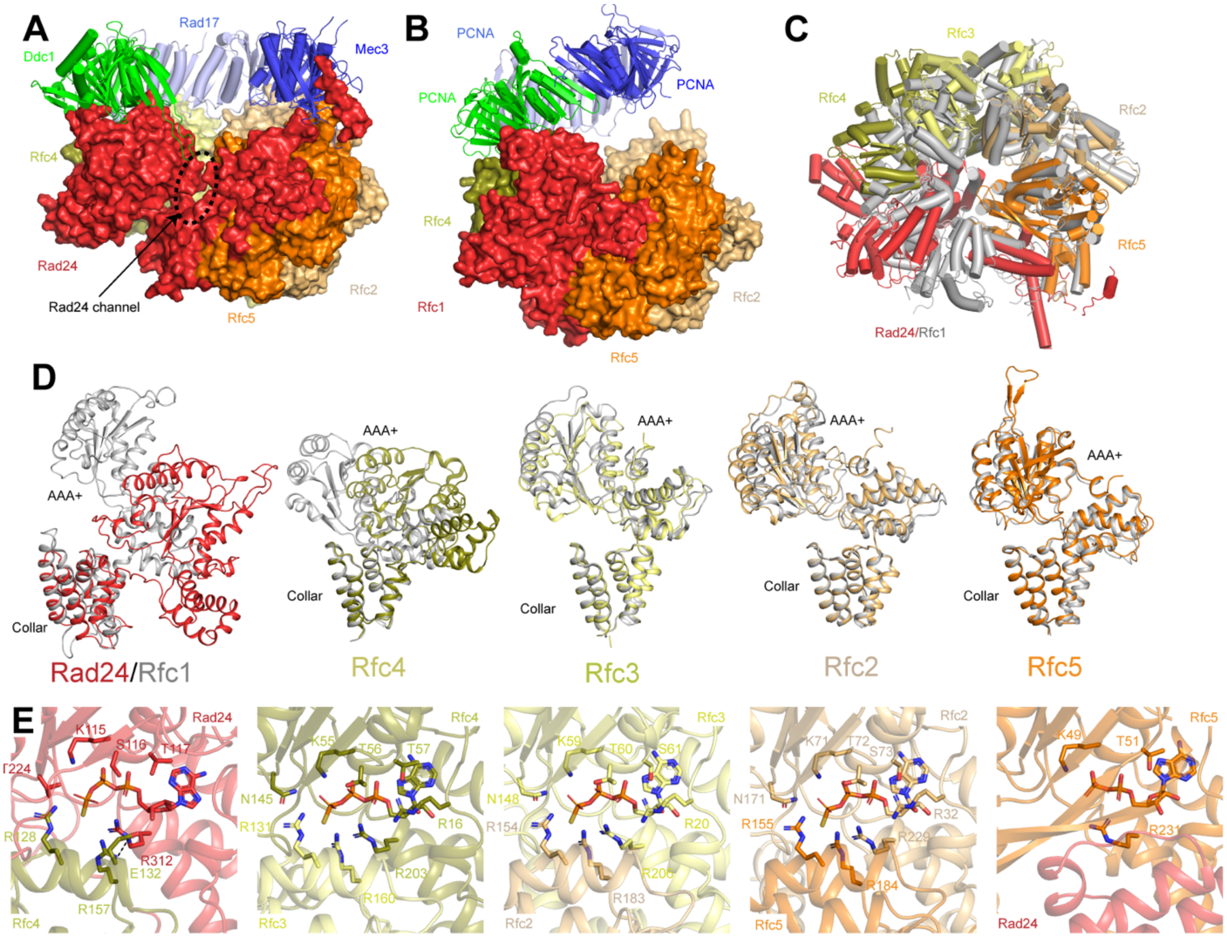


**Fig. S3. Comparison of Rad24-RFC:9-1-1 with RFC-PCNA. (A-B)** Structures of Rad24-RFC:9-1-1 (**A**) and RFC-PCNA (**B,** PDB:1SXJ) colored by subunit with Rad24-RFC / RFC in surface depiction and 9-1-1 / PCNA in ribbon depiction. The channel formed between the AAA+, collar and A' domains of Rad24 in **a** is highlighted by the dashed oval. **(C)** Superposition of Rad24-RFC (colored by domain) with RFC (grey). The models are aligned by the collar domains. **(D)** Superposition of Rad24-RFC subunits (colored by subunit) with RFC subunits (grey). The subunits are aligned by their collar domains. For Rad24 and Rfc1, the A' domains are removed for clarity. **e**, Nucleotide-binding sites in the AAA+ domains Rad24-RFC:9-1-1. Nucleotides and residues whose side chains coordinate the nucleotide are shown in sticks.

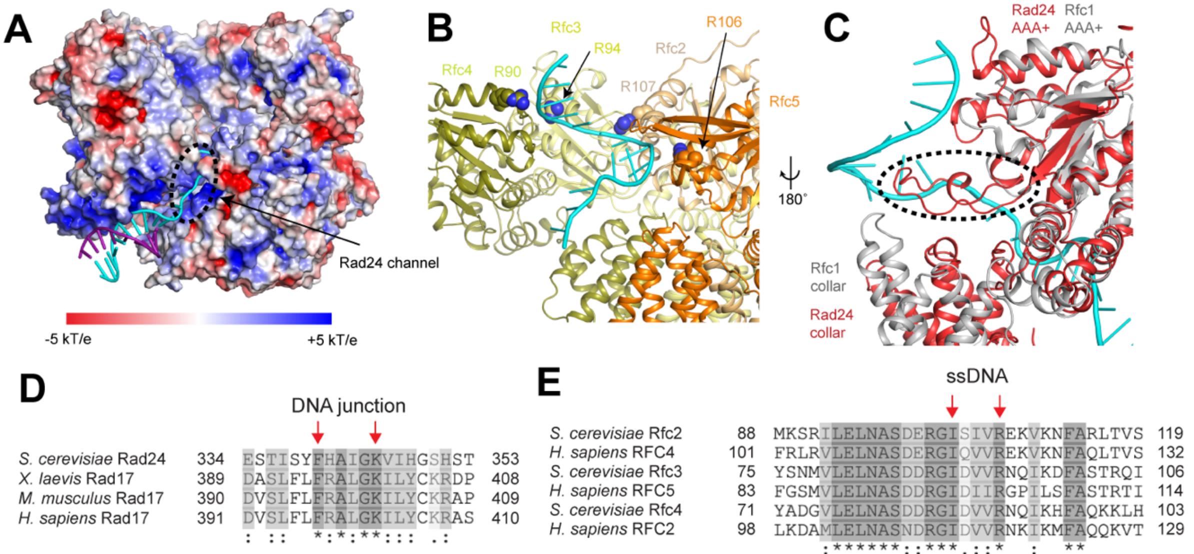


**Fig. S4. Coordination of DNA by Rad24-RFC:9-1-1. (A)** Rad24-RFC:9-1-1 surface colored by electrostatic surface potential. DNA is shown as cartoon and Rad24 channel is highlighted by the dashed oval. **(B)** Interactions between ssDNA backbone and conserved arginine residues in Rfc2-5 establish the spiral B-form like conformation of the ssDNA. Arginine side chains are shown as spheres and Rad24 is removed for clarity. **(C)** Superposition of Rad24 in open Rad24-RFC:9-1-1 (colored by subunit) with Rfc1 in RFC-PCNA (colored in grey, PDB:1SXJ), aligned by the AAA+ domains. Single-stranded DNA is shown as cartoon. Dashed oval highlights insertion in the AAA+ domain of Rad24 that occludes double-stranded DNA from occupying the central cavity of Rad24-RFC. Rfc2-5 are removed for clarity. **(D)** Alignment of residues that coordinate the 5' junction in *S. cerevisiae* Rad24 with Rad17 from *X. laevis*, *M. musculus,* and *H. sapiens*. Arrows denote conserved Phe340, which serves as the pin, and conserved Lys345, which coordinates the 5' phosphate. **(E)** Alignment of residues that coordinate the single-stranded DNA in *S. cerevisiae* Rfc2, Rfc3 and Rfc4 with RFC4, RFC5 and RFC2 from *H. sapiens*. Arrows denote conserved isoleucine and arginine residues that coordinate the DNA backbone.

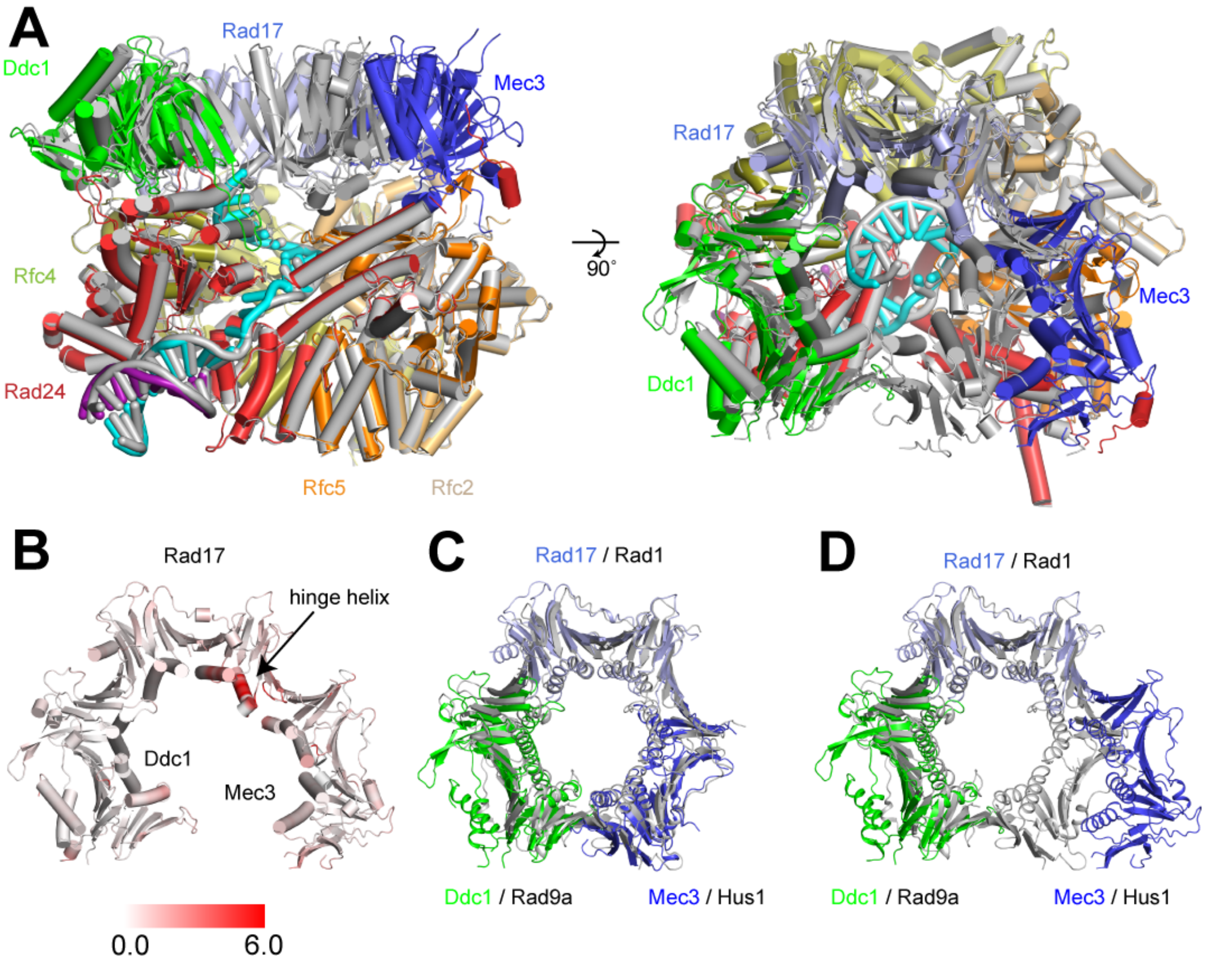


**Fig. S5. Comparison of open and closed states of Rad24-RFC:9-1-1. (A)** Superposition of open (colored by subunit) and closed states of Rad24-RFC:9-1-1 (colored in grey), shown as two views. **(B)** Structure of 9-1-1 in the open state of Rad24-RFC:9-1-1, colored by RMSD. RMSD is calculated on per-subunit basis between the open and closed states of Rad24-RFC:9-1-1. **(C,D)** Superposition of 9-1-1 clamp in Rad24-RFC:9-1-1 in closed (**C**) and open (**D**) states (colored by subunit) with a structure of human 9-1-1 clamp (colored in grey, PDB:3A1J).

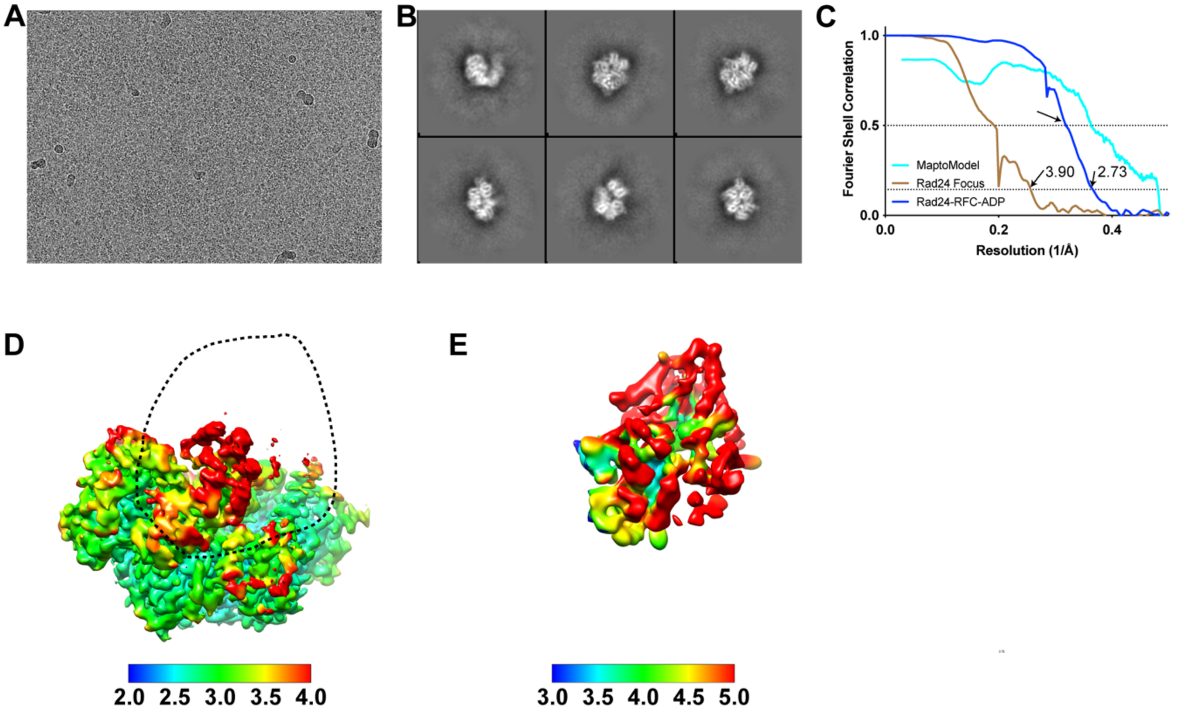


**Fig. 6. Validation of Rad24-RFC$_{ADP}$ structure. (A)** Representative cryo-EM image of vitrified Rad24-RFC in the presence of ADP. **(B)** Representative two-dimensional averages of Rad24-RFC$_{ADP}$. **(C)** Plot of Fourier shell correlations between two independent consensus half-maps (blue), two independent Rad24 focus refinement half-maps (brown), and the consensus map and model (cyan). **(D,E)** Cryo-EM density maps of consensus (**D**) and Rad24 focus refinement (**E**) maps colored by local resolution. Region included in the mask for the Rad24 AAA+ focused refinement (**E**) is denoted by the dashed line in **D**.

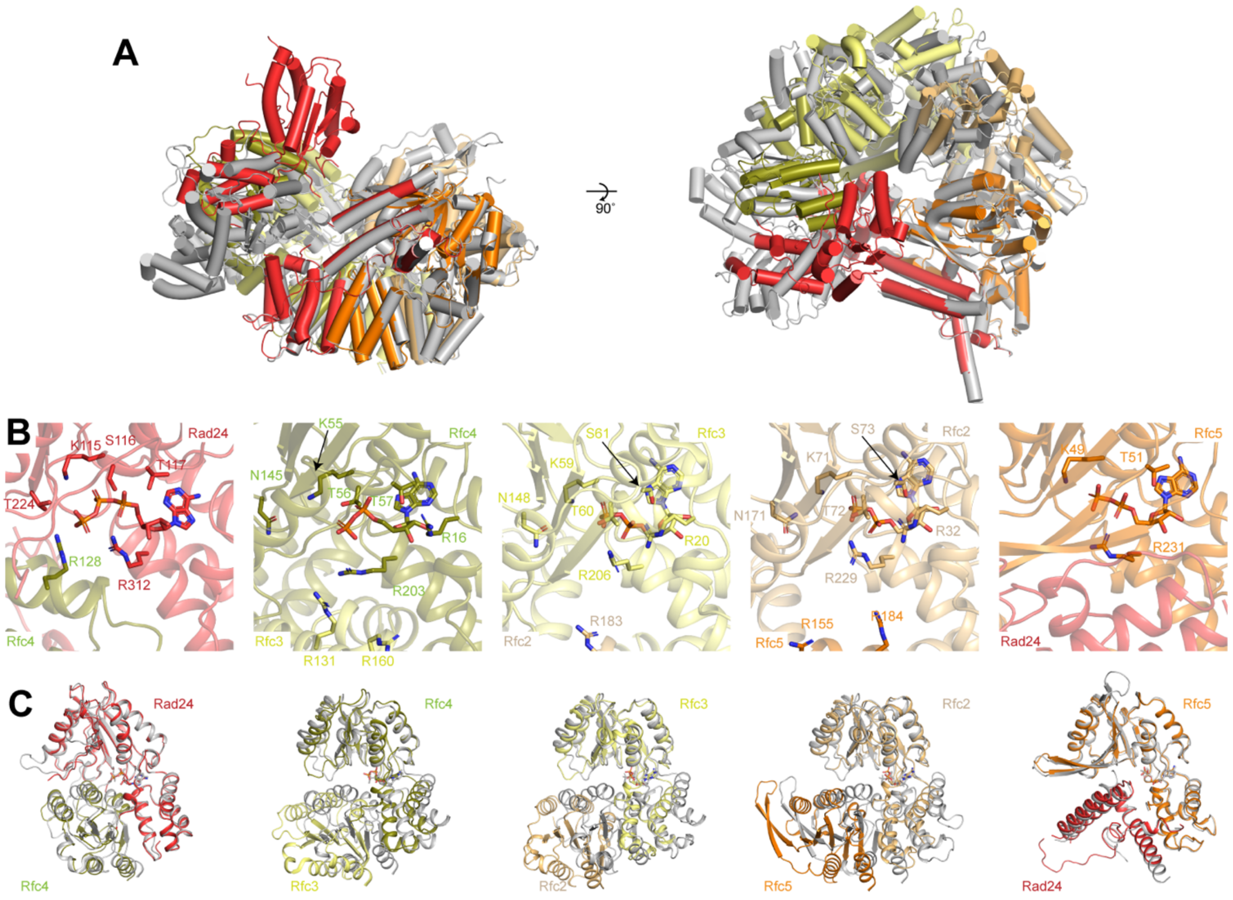


**Fig. S7. Conformational changes induced by ATP hydrolysis in Rad24-RFC.** a, Superposition of Rad24-RFC in Rad24-RFC$_{ADP}$ (colored by subunit) and Rad24-RFC:9-1-1 (colored in grey) shown in two views. b, Nucleotide-binding sites in the AAA+ domains of Rad24-RFC$_{ADP}$. Nucleotides are shown as sticks. c, Superposition of the AAA+ inter-domain domain interfaces in Rad24-RFC$_{ADP}$ (colored by subunit) and Rad24-RFC:9-1-1 (colored in grey). Nucleotides are shown as sticks.

**Table S1. Plasmids generated in this study.**

| Name | Insert |
|---|---|
| **pRS305G3-FLAG-Ddc1-Mec3++** | Ddc1 (synthetic ORF, N-terminal 3x-FLAG tag), Mec3 (synthetic ORF) |
| **pRS306G-Rad17++** | Rad17 (synthetic ORF) |
| **pRS305G-Rad24-FLAG++** | Rad24 (synthetic ORF, C-terminal 3x-FLAG tag) |

**Table S2. Summary of purification steps for individual proteins.**

| Protein | Tag | Purification steps |
|---|---|---|
| **9-1-1** | 3x FLAG at N-terminus of Ddc1 | 1. Anti-FLAG immunoprecipitation<br>2. Mono Q<br>3. Superdex 200 |
| **Rad24$^{Rfc}$** | 3x FLAG at C-terminus of Rad24 | 1. Anti-FLAG immunoprecipitation<br>2. Mono Q |
| **RPA** | None | 1. HiTrap Blue HP Chromatography<br>2. ssDNA affinity purification<br>3. Mono Q |

**Table S1. Yeast strains generated in this study.**

| Name | Genotype |
| --- | --- |
| **YJC3** | *MATa ade2-1 ura3-1 his3-11,15 trp1-1 leu2-3,112 can1-100 pep4::kanMX bar::hphNAT1 (hygromycinB) Gal-Gal4 (HIS3) Gal-Rfc2-Rfc3(TRP1) Gal-Rfc4-Rfc5 (URA3) Gal-Rad24 (LEU2)* |
| **YJC8** | *MATa ade2-1 ura3-1 his3-11,15 trp1-1 leu2-3,112 can1-100 pep4::kanMX bar::hphNAT1 (hygromycinB) Gal-Gal4 (HIS3) Gal-Rad17 (URA3) Gal-Mec3-Ddc2 (LEU2)* |

**Table S4. Cryo-EM data collection, refinement and validation statistics.**

| | Rad24-RFC-911 open (EMDB-xxxx) (PDB xxxx) | Mec3 focus refinement (EMDB-xxxx) (PDB xxxx) | Rad24-RFC-911 closed (EMDB-xxxx) (PDB xxxx) | Rad24-RFC-ADP (EMDB-xxxx) (PDB xxxx) | Rad24 AAA+ Focus refinement (EMDB-xxxx) (PDB xxxx) |
|---|---|---|---|---|---|
| **Data collection and processing** | | | | | |
| Magnification | 29,000x | 29,000x | 29,000x | 29,000x | 29,000x |
| Voltage (kV) | 300 keV | 300 keV | 300 keV | 300 keV | 300 keV |
| Electron exposure (e–/Å$^2$) | 66 | 66 | 66 | 66 | 66 |
| Defocus range (μm) | -0.5 to -2.0 | -0.5 to -2.0 | -0.5 to -2.0 | -0.5 to -2.0 | -0.5 to -2.0 |
| Pixel size (Å) | 0.813 | 0.813 | 0.813 | 0.813 | 0.813 |
| Symmetry imposed | C1 | C1 | C1 | C1 | C1 |
| Initial particle images (no.) | 4,117,022 | 4,117,022 | 4,117,022 | 4,432,412 | 4,432,412 |
| Final particle images (no.) | 938,420 | 938,420 | 237,512 | 183,334 | 81,834 |
| Map resolution (Å) | 2.20 | 2.43 | 2.72 | 2.73 | 3.90 |
| FSC threshold | 0.143 | 0.143 | 0.143 | 0.143 | 0.143 |
| Map resolution range (Å) | 200-2.20 | 200-2.43 | 200-2.72 | 200-2.73 | 200-3.90 |
| Density modification resolution (Å) FSC threshold | | | | 2.63<br>0.5 | 3.89<br>0.5 |
| **Refinement** | | | | | |
| Initial model used (PDB code) | 1SXJ, 3A1J | | Open state | Open state | |
| Model resolution (Å) | 2.20 | | 2.87 | 2.75 | |
| FSC threshold | 0.5 | | 0.5 | 0.5 | |
| Model resolution range (Å) | 200-2.20 | | 200-2.87 | 200-2.75 | |
| Model composition | | | | | |
| Non-hydrogen atoms | 23,622 | | 22,813 | 14,249 | |
| Protein residues | 2802 | | 2753 | 1774 | |
| Ligands | 9 | | 9 | 6 | |
| *B* factors (Å$^2$) | | | | | |
| Protein (mean) | 78.5 | | 87.0 | 83.8 | |
| Ligand (mean) | 11.1 | | 39.0 | 89.4 | |
| R.m.s. deviations | | | | | |
| Bond lengths (Å) | 0.002 | | 0.002 | 0.002 | |
| Bond angles (°) | 0.443 | | 0.446 | 0.509 | |
| Validation | | | | | |
| MolProbity score | 1.08 | | 1.32 | 1.17 | |
| Clashscore | 2.86 | | 4.07 | 3.87 | |
| Poor rotamers (%) | 0.71 | | 0.00 | 0.00 | |
| Ramachandran plot | | | | | |
| Favored (%) | 98.55 | | 97.34 | 98.05 | |
| Allowed (%) | 1.45 | | 2.62 | 1.95 | |
| Disallowed (%) | 0.00 | | 0.04 | 0.05 | |
